# Concurrent visual working memory bias in sequential integration of approximate number

**DOI:** 10.1101/2020.07.16.191445

**Authors:** Zhiqi Kang, Bernhard Spitzer

## Abstract

Previous work has shown bidirectional crosstalk between Working Memory (WM) and perception such that the contents of WM can alter concurrent percepts and vice versa. Here, we examine WM-perception interactions in a new task setting. Participants judged the proportion of colored dots in a stream of visual displays while concurrently holding location- and color information in memory. Spatiotemporally resolved psychometrics disclosed a modulation of perceptual sensitivity consistent with a bias of visual spatial attention towards the memorized location. However, this effect was short-lived, suggesting that the visuospatial WM information was rapidly deprioritized during processing of new perceptual information. Independently, we observed robust bidirectional biases of categorical color judgments, in that perceptual decisions and mnemonic reports were attracted to each other. These biases occurred without reductions in overall perceptual sensitivity compared to control conditions without a concurrent WM load. The results conceptually replicate and extend previous findings in visual search and suggest that crosstalk between WM and perception can arise at multiple levels, from sensory-perceptual to decisional processing.

## Introduction

What is on our mind can affect how we perceive the physical world. In line with this intuition, laboratory experiments have shown that the contents of working memory (WM) can impact on performance in intervening perceptual tasks. For instance, when actively maintaining a green item in WM while searching for a target shape, response times can be slowed when a green distractor is present in the search array^1,2^, indicating attentional capture by perceptual input that matches the concurrent WM content^1–4^. The attention-guiding effect of WM has been linked to a view that WM information can be maintained in different functional states^1,5,6^. According to this view, WM contents guide attention while they are prioritized for immediate goal-directed use. WM contents that are not currently prioritized can be maintained for prospective use on later occasion, but evidence suggests that they do not bias attention in concurrent perceptual tasks^1,7^.

Beyond attentional guidance, several studies have shown that concurrent WM information can even bias the very appearance of intervening stimuli. For instance, the perceived orientation or motion of a stimulus was found to be repulsed away from that of a concurrently maintained stimulus^8–10^. One recent study instead suggested that sensory representations can be attracted towards concurrent memories^11^. However, crosstalk between WM and perception has also been found to occur the other way round, such that intervening stimuli can bias later recall of concurrently maintained information^9–12^. The typical pattern in this type of finding is that memory reports are attracted to (rather than repulsed away from) intervening stimuli^10,11,13^.

In explaining WM-perception biases, previous studies have assumed more or less direct alterations in the representation of the perceptual and/or mnemonic information^10,11,13^. WM representations may drift or shift^14,15^ towards concurrent percepts and become noisier^10^. And vice versa, it has been suggested that concurrent WM contents may directly alter the sensory tuning to new input^11^. However, from a decision-theoretic perspective, biases in behavioral reporting may also arise in post-perceptual evaluation of otherwise unchanged sensory and/or mnemonic representations^16^. Recent work on trial history effects, for instance, found attraction biases in subjective reporting, but repulsive biases in more direct psychophysical measures of stimulus appearance^17^. Attractive WM-perception biases might also arise if concurrent contents induced a tendency for congruent reporting, and/or by partial confusion of remembered and just-experienced information. Such bias could be characterized as a shift in response criteria^16^ or, in a sequential sampling framework^18^, as an offset of decisional evidence accumulation towards the response category that is associated with the concurrent information.

The attention-guiding effects of WM have typically been demonstrated in unrelated tasks that demand rapid responding to a single display (e.g., a visual search array). Little is known if and for how long attentional guidance would persist beyond the timescale of these tasks, for instance when the to-be-judged information is spread out in time. Bidirectional crosstalk between perceptual and mnemonic reports, on the other hand, can only be observed when the perceptual- and the WM task share the same relevant stimulus dimension (e.g., judging orientation while remembering another orientation). To what extent attraction- or repulsion biases in the latter kind of tasks depend on the functional state of the WM information is not clear yet.

Here, we examined WM-perception interactions in a sequential integration task that required participants to judge visuospatial numerosity information in a stream of random dots displays while concurrently remembering the location and color of a WM sample for later report. Color was independently task-relevant also in the decision task, which enabled us to examine spatial attentional bias along with bidirectional biases of perceptual and mnemonic decisions. We found evidence for a bias of visual spatial attention towards the to-be-memorized location which, however, dissipated quickly after the onset of the decision stream. Bidirectional bias of perceptual and mnemonic reports instead emerged in form of additive choice biases and categorical misreporting, in line with a post-perceptual locus of interference.

## Results

Participants (n=68) were asked to remember the color and location of a WM sample stimulus (Fig. 1a, *left*) while deciding whether an intervening stream of six random dots displays contained relatively more blue or red dots (Fig. 1a, *middle*). After choice, subjects were asked to reproduce the color and spatial location of the WM sample from memory (Fig. 1a, *right*). Thus, the WM task involved maintaining both categorical information about a feature that was task-relevant also in the decision task (red/blue) and high-precision information about stimulus location which was not to be judged or reported in the intermittent task. Decision- and WM reports were entered with different hands and using distinct button operation procedures (select or move and toggle, see *Methods*) to avoid motor response confusion between the two tasks.

**Figure 1.**
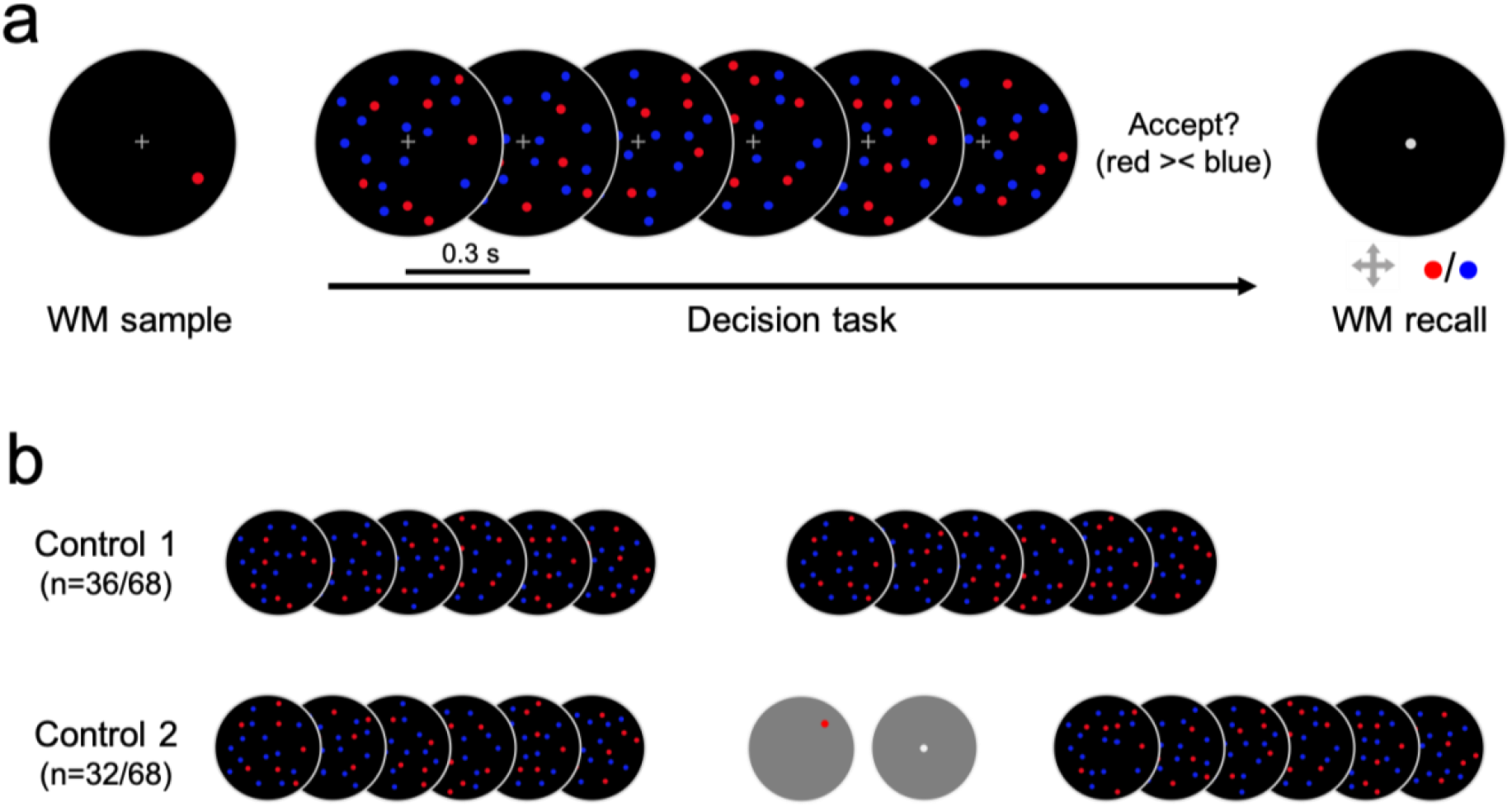
Experimental paradigm. **a,** Schematic outline of a trial in the WM interference condition. *Left*, the to-be-maintained WM sample was a single dot (red or blue) presented at a random position on an invisible circular path around fixation. *Middle*, During WM maintenance, a stream of 6 random dots displays was presented. Each display contained 20 dots, a variable number of which was blue, the others red. Participants were asked to evaluate whether the stream contained relatively more blue or red dots (see *Methods* for details). *Right*, both the color and location of the WM sample were to be reproduced from memory at the end of the trial. **b,** Within-subjects control conditions. In Control 1, the WM task elements were omitted. In Control 2, the WM- and decision task elements were rearranged such that the two tasks were not concurrent.

We asked if the concurrently memorized WM sample would alter visuospatial processing of the decision stream. First, we mapped the participants’ overall spatial weighting through logistic regression of perceptual reports against the trial-by-trial varying color value in each pixel of the decision displays. The spatial weights were compared against those predicted by an individually fitted null model as unbiased baseline (see *Methods*). Figure 2a illustrates the grand mean weighting map, collapsed across all experimental conditions (WM and controls) and display positions (1-6). Participants were most sensitive towards information in a central vertical region of the display (p<0.05, FDR-corrected across pixels) while giving less weight to the lateral periphery, especially at the right-hand side of the display area (p<0.05, FDR-corrected). This pattern seems to differ from classic findings of heightened sensitivity along the horizontal meridian in studies of low-level visual acuity^19–21^. However, the pattern we observed could be related to the notion of an anisotropy of perceived space, by which subjects estimate spatially spread out magnitudes like line length^22^ and numerosity^23^ to be larger along the vertical axis. We anticipate this incidental observation to be of interest for vision- and numerical cognition scientists using similar stimuli. To verify if the weighting pattern was stable throughout the stream sequence, we examined the pairwise spatial correlations between the weight distributions in each of the six displays. All of the 15 pairwise correlations were positive (mean R=0.65, min 0.52, max 0.78) and remained all positive when subtracting the mean correlations obtained after randomly rotating each map (1000 iterations; which excludes center-bias; mean R=0.32, min 0.17, max 0.44). Our method thus proved efficient in disclosing a robust and stable spatial weighting pattern in the stream task.

**Figure 2.**
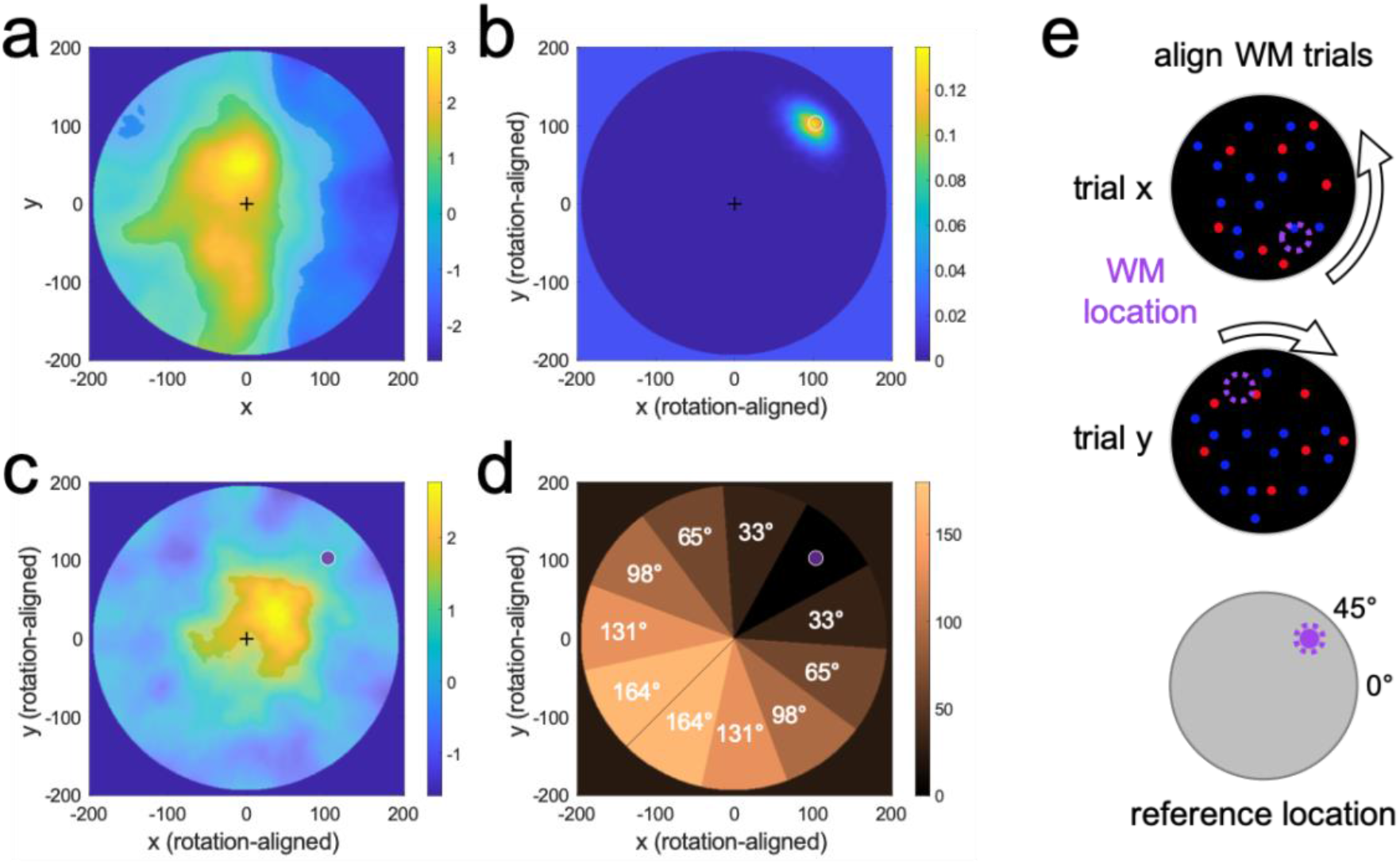
Spatial weighting analysis. **a,** Spatial weighting of the decision displays before rotational alignment, collapsed across all trials (WM and control conditions) and displays (1-6). Positive values indicate overweighting, negative values underweighting, relative to an unbiased observer model fitted to each individual (see *Methods*). Transparent mask indicates significant regional over- or underweighting (p<0.05, two-tailed, FDR-corrected across pixels). **b,** Spatial distribution of WM sample positions reported on WM recall (cf. Fig. 1a, *right*) after rotational alignment (cf. *e*), aggregated across all participants. White circle indicates true position (rotation-aligned) of the WM sample. **c,** Spatial weighting on WM trials after rotational alignment (cf. *e*), same conventions as in b. Purple dot indicates (rotation-aligned) location of the concurrently maintained WM sample. **d,** Pie masks for angular tuning analysis. Spatial weights within each segment were averaged and examined as a function of the absolute angular distance from the WM-sample (see Fig. 3b below for results). **e,** Rotational alignment of trials. Displays were rotated offline such that the trial-specific WM sample positions matched the same (virtual) reference location (arbitrarily set to 45°, cf. purple markers).

Comparing the overall weighting pattern in Fig. 2a between trials with and without a concurrent WM load, we found no significant differences (no pixels p<0.05, FDR-corrected for multiple comparisons). However, our main question was whether on WM trials, the allocation of processing gain in visual space was attracted towards (or repulsed away from) the to-be-memorized WM sample location. To this end, we offline rotated the circular displays from all trials such that they were all aligned to the same WM sample position (Fig. 2e). We arbitrarily set this common reference position to 45° for illustration purposes. The weighting map computed from the thus aligned displays, averaged across displays (1-6), indeed showed a concentration of sensitivity towards the WM sample location (Fig. 2c, p<0.05, FDR-corrected across pixels). In other words, the concurrent WM sample directly modulated the gain of perceptual processing in visual space. This effect manifested as a moderate shift of sensitivity away from fixation in the direction of the WM sample, rather than a concentration at its actual physical position (which was more peripheral and reproduced with high accuracy on later recall, cf. Fig. 2b). Unlike the overall spatial weighting pattern (cf. Fig. 2a), the WM-related spatial bias (Fig. 2c) appeared not to be stable over time (mean inter-display correlation after rotational alignment, R=−0.03, min −0.30, max 0.17).

For further analysis, we split the display area into equal-sized pie segments, with a target segment centered around the WM sample location at 45° (Fig. 2d). Averaging weight within segments allowed us to examine the “tuning” of spatial weighting, in terms of its mean angular distance from the WM sample location. Figure 3b illustrates the angular tuning separately for each of the six displays in the decision stream. Statistical analysis of the mean weight in the WM sample segment compared to the remaining segments showed a significant effect in the first stream display [display 1; t(67)=4.19, p<0.001]. This angular tuning was even discernable directly in the weighting map of display 1, with significantly increased sensitivity (p<0.05, FDR-corrected across pixels) along the trajectory between fixation and WM sample location (leftmost in Fig. 3a). However, no such effect was evident in any of the subsequent displays (2-6), neither in pixel-level weighting maps (no pixels p<0.05, FDR-corrected) nor in terms of angular tuning [all t(67)<1.66, p’s >0.10]. Bayes factor analysis indicated strong evidence for an angular tuning effect in display 1 (BF01=0.004), but anecdotal to moderate evidence for its absence in displays 2-6 (BF01 ranging from 2.06 to 4.85). For further validation, we pseudo-aligned the trials from control conditions (cf. Fig. 1b) to the same WM sample locations as the WM trials (i.e., *as if* each WM sample had been presented on a control trial, too). This control analysis showed no (pseudo-) angular tuning for any of the six displays (yellow in Fig. 3b, all p’s >0.05).

**Figure 3.**
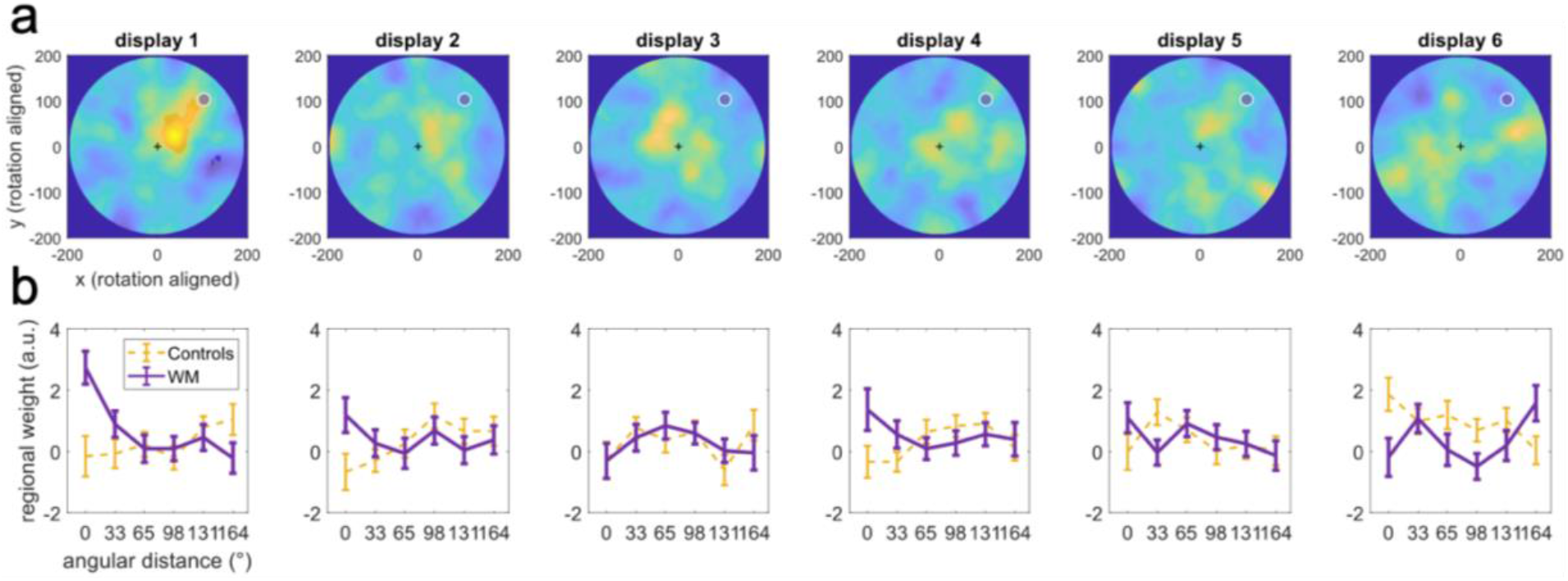
Regional weight concentration – time course. **a,** spatial weighting on WM trials after rotational alignment as in Fig. 2c but shown separately for each of the six displays in the decision stream. A significant regional gain concentration (p<0.05, two-tailed, FDR-corrected, indicated by transparent mask) was observed in display 1 only. **b,** *Purple:* angular tuning of spatial weighting in terms of mean angular distance from the WM sample (cf. Fig. 2d). Significant tuning was evident exclusively in display 1. Yellow curves show analogue analysis of control trials (pooled over control conditions 1 & 2) pseudo-aligned to the same WM-locations as the WM-trials.

To verify if the spatial weighting bias was related to WM processing, we median-split each participant’s trial data according to the precision of location reports on subsequent WM recall. We observed significant angular tuning in display 1 on high-precision WM trials [t(67)=4.61, p<0.001, BF01=0.001] but not on low-precision WM trials [t(67)=1.21, p=0.23, BF01=3.74]. Thus, the presentation of a WM sample alone was not sufficient to explain the weighting shift on WM trials. This finding, together with the relatively small radius of the shift, also renders the effect in display 1 less likely to have resulted from reflexive saccades to the WM sample (but see Discussion for a potential role of microsaccadic eye activity in covert spatial attention).

Together, visuospatial processing was temporarily biased towards the location of the WM sample, but this bias was ephemeral and dissipated quickly with new perceptual decision information. Visual processing of the remainder of the stream appeared unaffected by the concurrently maintained spatial WM information. The pattern suggests that the focus of spatial attention swiftly moved away from the WM sample when the decision task stream commenced.

Our analysis has thus far focused on interactions with the spatial information in the WM sample. We next asked to what extent the concurrently memorized categorical color information, which was also remembered with high accuracy (mean percentage correct 93.1 %, SE=0.006), may have biased judgments of the intervening decision stream. To this end, we examined psychometric weighting of the blue-red composition of the stream displays using conventional reverse correlation analysis^24–26^. The weighting functions in Figure 4a indicate the proportion of blue>red choices as a function of the relative blue-red dot count in a display. Descriptively, the mean weighting functions were near-linear and appeared parallelly shifted towards the color of the WM sample (blue or red), indicating an additive decision bias, compared to control conditions without a concurrent WM load (Fig. 4a). A repeated measures ANOVA with the factors WM condition (blue, red, control) and dot count (11 levels) showed a main effect of WM condition [F(1.96,131.62)=29.97, p<0.001] but no interaction with dot count [F(12.52,838.54)=1.158, p=0.31], consistent with a parallel displacement of the psychometric function [main effect of dot count: F(6.55,438.77)=1041.44, p<0.001]. We next fitted a logistic choice model (see *Methods*) to independently quantify the psychophysical sensitivity (slope) towards the stream displays’ color composition, and the strength of an additive choice bias (intercept) associated with maintaining a blue or red WM sample, respectively.

**Figure 4.**
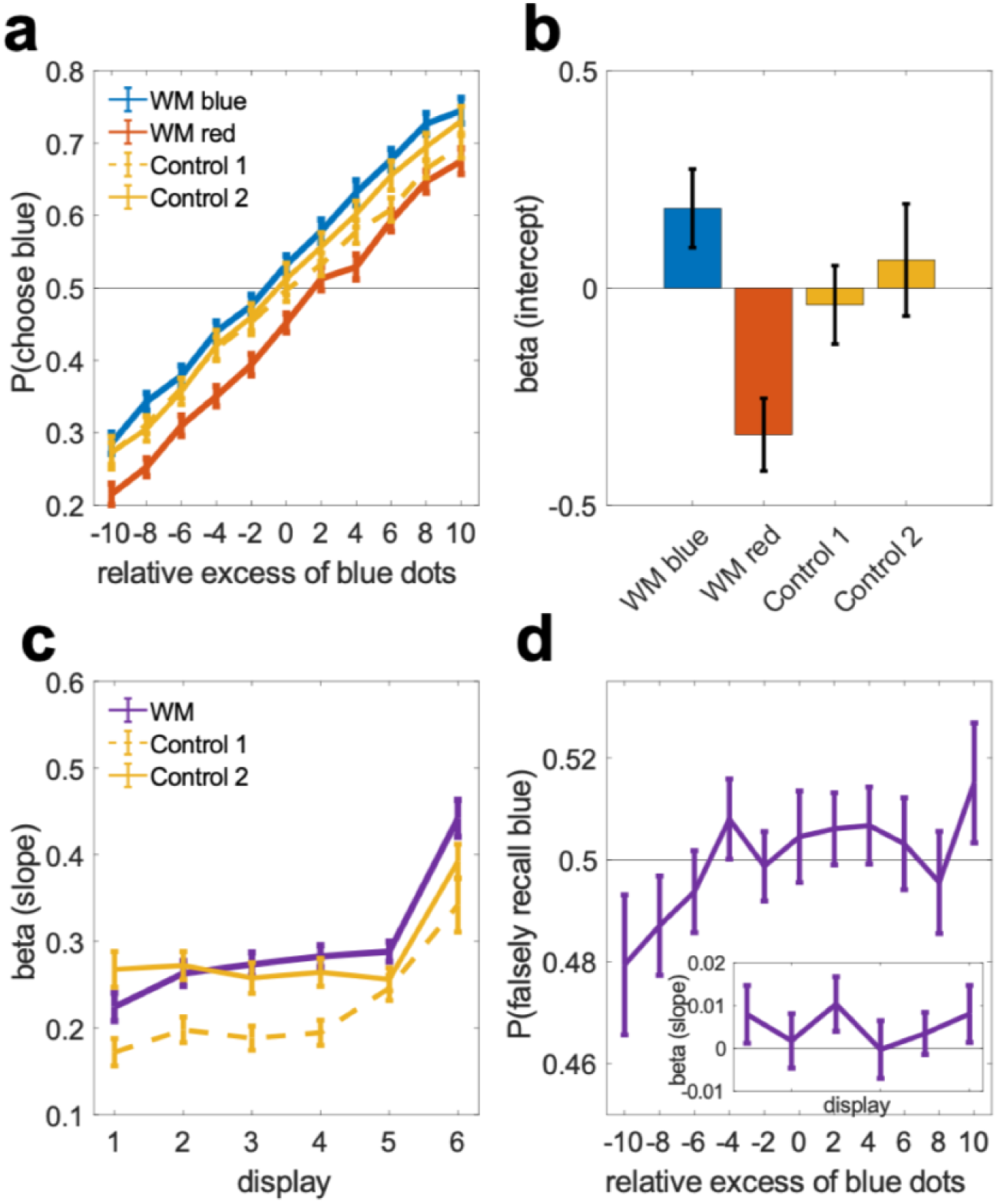
Bidirectional biases of WM and perceptual decisions – color information. **a,** psychometric weighting functions averaged over the six displays in the decision task, plotted separately for trials where the concurrently maintained WM sample was red or blue, and for control conditions without a concurrent WM task (cf. Fig. 1). **b,** bias terms (intercepts) derived from logistic regression of choice. **c,** choice sensitivity (slopes) to the red-blue dot composition in each of the six displays in the decision stream. **d,** Memory recall bias. Probability of (erroneous) blue/red report in WM recall as a function of the blue-red composition of the intervening decision displays. Small inset plot shows the time-course of this relation over the six stream displays in terms of logistic regression coefficients. Error bars in all panels show standard error of the mean.

Statistical analysis of the model coefficients corroborated a significant choice bias towards the WM sample color [Fig. 4b, intercepts; WM blue: t(67)=2.64, p<0.02, WM red: t(67)=5.11, p<0.001, paired t-tests against pooled control conditions]. We confirmed that this additive effect was associated with active WM maintenance and not merely with the presentation of a WM sample: the magnitude of the choice bias (WM blue – WM red; cf. Fig.4b) was significantly enhanced when restricting the analysis to those trials in which the WM color was subsequently recalled correctly [0.598 vs. 0.521, t(67)=4.3877, p<0.001]. Repeating the analysis with subjects’ color recall reports (instead of physical WM sample color) as bias terms descriptively increased the bias (0.561 vs. 0.521), but this difference failed to reach significance [t(67)= 1.538, p=0.13; see below for targeted analysis of reporting-level bias]. Interestingly, when median splitting trials according to the precision of WM *location* recall instead, the choice bias did not differ [0.572 vs. 0.501, t(67)=0.54, p=0.59] and was robustly present in both split sets [both t(67)>4.5, both p’s<0.001]. Thus, the color bias was observed regardless of whether or not a spatial bias was evident on the same trials (cf. angular tuning analysis above), indicating a degree of independence between the two effects.

Turning to sensitivity (Fig. 4c, slopes), we observed an overall recency effect^27,28^, with heightened sensitivity towards the end of the decision stream [main effect of display position 1-6: F(2.67,179.07)=43.94, p<0.001], which may reflect memory loss or “leakage” of early presented decision information^28,29^. The recency effect was slightly stronger on WM trials, with a steeper time course compared to pooled controls [difference in slope of increase: t(67)= 2.45, p<0.02], indicating that the concurrent WM load may have interfered with the mnemonic demands of the stream task. Interestingly, we found no reduction in overall sensitivity during WM maintenance compared to control conditions without a concurrent WM load. In fact, in one of the control conditions (Control 1), sensitivity was even significantly lower than on WM trials [Fig. 4c; main effect of condition, F(1,35)=55.37, p<0.001]. This unexpected observation could be attributable to procedural aspects (decision trials occurred in faster succession in the Control 1 condition). In our second control condition (Control 2), which controlled for these aspects (cf. Fig. 1b), overall sensitivity was statistically indistinguishable from that on WM trials [F(1,31)=0.734, p=0.40)]. To summarize, concurrent WM maintenance robustly biased choices and slightly increased recency effects, but did not impair overall perceptual-discriminative acuity.

We analogously examined whether information in the decision stream would bias subsequent recall of the WM sample color (Fig. 4d). A logistic regression of WM color reports (blue/red) against the relative blue-red dot count in the intermittent stream displays showed a significant positive effect [t(67)=2.14, p=0.036, t-test of pooled slope coefficients against zero]. Thus, the more blue (or red) dots were contained in a stream display, the more likely the WM sample was erroneously recalled as blue (or red). The time course of this effect showed no significant variation across the six stream displays [inset in Fig. 4d; F(5,402)<1], suggesting that the bias was not only driven by e.g., early displays that occurred in temporal proximity to the WM sample.

Does this WM bias reflect a direct distortion of color memory by sensory input and/or a crosstalk with post-perceptual evaluations in the decision task? In further analysis, we regressed the WM color reports against the choice residuals from the psychometric decision analysis (cf. Fig. 4b-c; *Methods*, Eq. 2). Thus, we tested if WM reports were even biased by endogenous choice variability in the stream task that is unexplained by any stimulus information (neither in WM-nor decision samples). This analysis showed a positive effect [t(67)= 4.68, p<0.001], suggesting crosstalk on the level of decisional evaluation and/or - reporting. Finally, we also directly correlated the residuals in WM- and decision reports after applying the full psychometric model (*Methods*, Eq. 2) to either. The full model perfectly separated WM reports in the three highest performing participants. In the remaining subjects, we found a significant positive correlation between the residuals in WM- and decision reporting [mean R: 0.04, SE=0.008, t(64)=4.79, p<0.001]. We note again that our experimental setup rendered such crosstalk unlikely to arise from simple motor response- or button confusion (see *Methods*).

We explored yet another task aspect in our experiment: one group of participants was asked to judge if the stream contained more blue (or red) dots on average (“averaging” condition) which mirrors the typical task requirement in perceptual choice experiments. Another group was asked to decide if they were willing to receive the value of one randomly drawn sample from the just presented stream (“gambling” condition), which mimics the scenario of an economic “risky” choice task. Normatively, observers in both these conditions should behave identically in order to maximize long-term returns. However, we hypothesized that the gambling scenario could promote a more discretized representation of the individual stream displays and thus load more strongly on WM storage processes^30^, whereas the averaging task may promote more continuous updating of a running decision variable^18,28,31^, potentially posing a lower WM load. We therefore anticipated the gambling task to interfere more strongly with the concurrent WM task.

We observed no differences in spatial weighting (cf. Fig. 2a) or angular tuning (cf. Fig. 2c) between the two task variants (no pixels p<0.05, FDR-corrected). We also found no reliable differences in color bias (Fig. 5a), neither in terms of its magnitude [t(66)=1.79, p=0.08, two-sample t-test] nor direction [t(66)=0.54, p=0.59]. Overall choice sensitivity (Fig. 5b) was significantly lower in gambling than in averaging [t(66)=2.64, p=0.01] which replicates earlier findings^32^. Furthermore, the analysis showed a robust WM-induced enhancement of recency in the gambling group [t(32)=3.23, p=0.003, difference in slope of increase across displays between WM- and control conditions] but not in the averaging group [t(34)=0.74, p=0.47]. However, the between-group difference was not significant [t(66)=1.33, p=0.19]. Finally, the difference in overall sensitivity between WM- and control trials did not differ between groups [t(66)=1.16, p=0.25].

**Figure 5.**
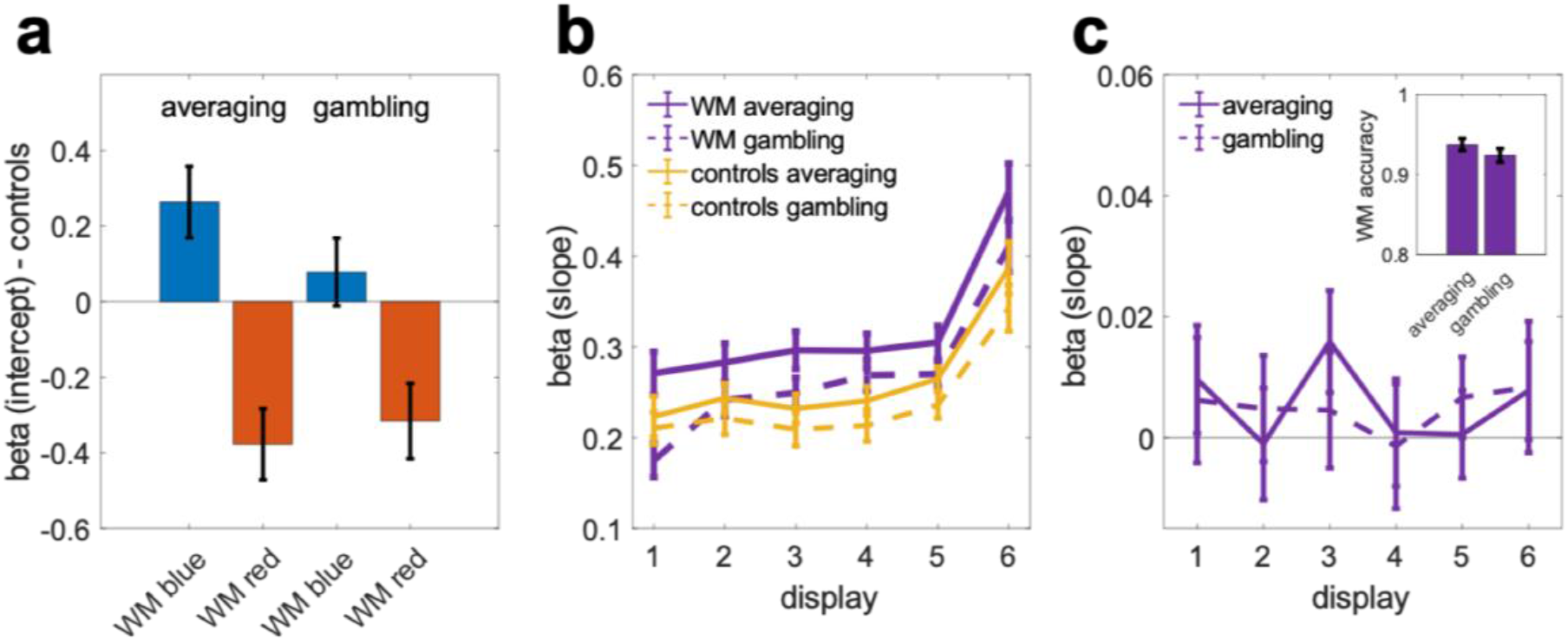
Averaging or gambling – color information. **a,** WM-induced choice bias relative to pooled control conditions (cf. Fig. 4b), plotted separately for “averaging” (left bars, n=35) and “gambling” variants (right bars, n=33) of the decision task. **b,** choice sensitivity (slopes) for each of the six displays in the decision stream (cf. Fig. 4c), plotted separately for the averaging and gambling tasks. Yellow curves show pooled control conditions. **c,** WM recall bias (cf. inset in Fig. 4d) plotted separately for the averaging and gambling conditions. Small inset plot shows proportion correct color recall. Error bars in all panels show standard error of the mean.

Examining the bias in subsequent WM-color recall (cf. Fig. 4d) separately for the averaging- and gambling groups, we found no difference between the two [Fig. 5c, t(66)=0.15, p=0.88; two-sample t-test of regression coefficients pooled over displays]. Overall WM color recall appeared numerically slightly less accurate after gambling (0.92) than after averaging (0.94, inset in Fig. 5c) but the difference was not significant [t(66)=1.20, p=0.23, two-sample t-test]. Lastly, we asked if WM location recall (cf. Fig. 2b) differed between the two variants of the intermittent decision task. We found that participants in the gambling condition tended to report the WM-sample locations less precisely (Fig. 6). Together, we found weak tendencies for the risky choice task to interfere slightly more with concurrent WM than the averaging task, but otherwise no difference in multiplicative (spatial) and/or additive (color) biases between the two task variants.

**Figure 6.**
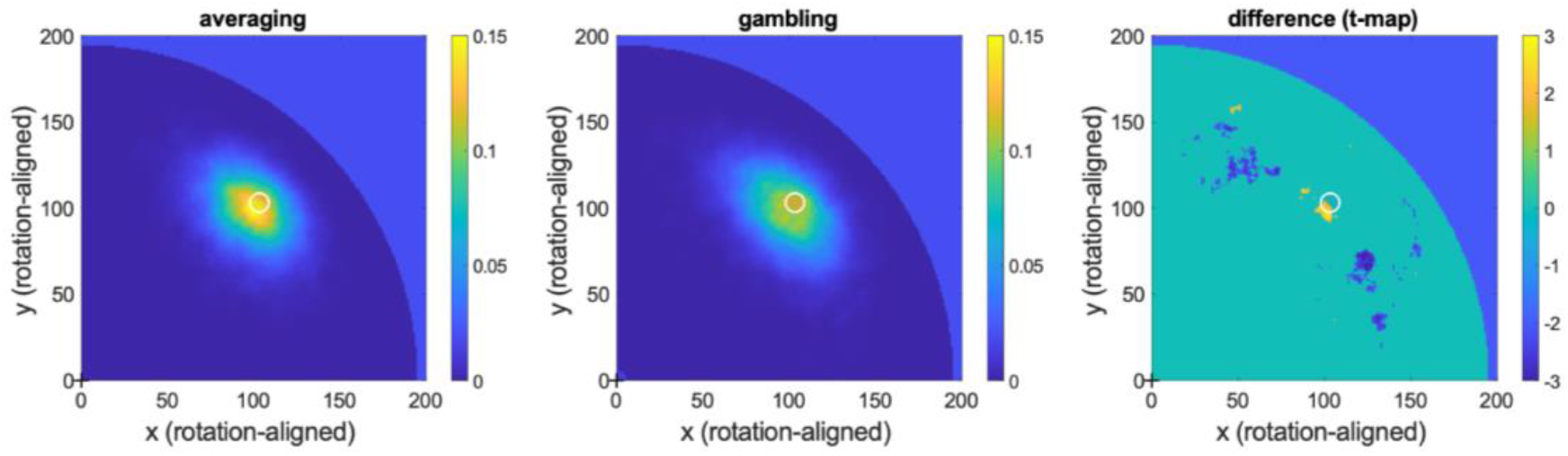
Averaging or gambling – spatial WM precision. Spatial distribution of WM positions reported on WM recall (rotation-aligned as in Fig. 2b), shown separately for the averaging- (*left*) and gambling (*middle*) variants of the intermittent decision task. *Right* panel shows statistical map of the difference between the two, thresholded at p<0.05 (two-tailed, uncorrected). White circles indicate true original position (rotation-aligned) of the WM sample. Participants in the averaging condition tended to report more locations near to the target and fewer locations afar from it, compared to the gambling condition.

## Discussion

To summarize our findings, we report an array of WM-perception interactions, including (i) a short-lived shift of perceptual sensitivity towards the just-encoded location of the WM item in visual space, (ii) a bidirectional bias to categorically (mis-)judge decision information according to the concurrent WM information and vice versa, and (iii) mild WM interference with the mnemonic demands of the decision task, in terms of enhanced recency and reduced spatial recall precision. Alongside this multifaceted crosstalk between WM and perception, we observed no general reduction in perceptual sensitivity.

Our sequential integration task enabled us to track spatial attentional bias in a time-resolved manner. The finding of a modulation of local visuospatial sensitivity conceptually replicates and extends previous findings in spatial WM^33^ and in visual search speed^1–4^. However, we found this effect to be short-lived, indicating that attentional guidance adapted rapidly to momentary task demands. From a state-based WM perspective^1,6^, the focus of attention appeared to swiftly move away from the WM sample location when the decision stream commenced. Participants nevertheless reproduced the WM location accurately on later recall, in line with previous evidence that temporary withdrawal of attention does not necessarily disrupt memory performance^34–36^. An interesting question for future work is how the present patterns in visuospatial weighting may relate to physiological indices of covert spatial attention e.g., in alpha-band EEG^37^ or microsaccadic eye activity^38^.

The time course of spatial bias reported here seemingly contrasts with recent findings of WM-induced attentional biases that persisted throughout several successive search- or choice displays after presentation of a WM sample^8,39^. However, in these previous studies, each display was to be judged separately before succeeding to the next, which may have enabled participants to periodically refocus on the WM information. Our challenging sequential integration task likely precluded such attentional reorienting to the WM content^see also 40^. Consistently, overall psychometric sensitivity in our task was unimpaired by the concurrent WM load^see also 41^. We found the spatial weighting bias to be most pronounced when the WM sample location was subsequently recalled with high precision, indicating a relation to memory-relevant processes. However, based on the present data, we cannot distinguish whether the transient bias was driven by the encoding- or the maintenance requirement of our WM task. Future studies utilizing e.g., retrospective cueing^42,43^ can help to shed light on this open question.

After prolonged, presumably unprioritized storage, the concurrent WM information still showed a marked influence on choice behavior in terms of a robust bias in categorical color judgments. Unlike the modulation of spatial sensitivity, the effect of WM color was additive in nature, i.e., it emerged independent of the very stimulus information on display^16^. One potential explanation for this effect could be in terms of selective visual processing. The WM sample may have led participants to prioritize those dots in the decision displays that matched the memorized color. Such prioritization could have occurred inadvertently, or participants might have used it as a deliberate memory aid (but see^4^ for arguments against strategic factors in WM-perception interactions). If selective processing had led to overestimation of relative numerosity^cf^. ^44,45^, and invariantly so across the entire stimulus range (red-blue), an additive bias might emerge. Psychometric modeling does not afford to infer the precise time point(s) when such effect may have arisen and whether it occurred, for instance, during the early and/or later periods of the decision stream. However, considering the high attentional and mnemonic demands of our stream task, and given the absence of spatial WM bias on later displays, it appears unlikely that the WM color would have guided visual processing for long, alongside the accumulating, immediately task-relevant information in the decision stream.

Another, simpler explanation is that the choice bias may have arisen on the level of decisional categorization. The categorical color information in the WM sample may have induced a baseline shift from the outset of evidence accumulation in the decision task. A similar account^see also 46^ has been proposed to explain previous findings that WM-matching stimuli seem to enter visual awareness more easily after interocular suppression^47,48,but see 49^. In our sequential integration task, an offset in evidence accumulation can give rise to a choice bias that is independent of the physical stimulus input, as was observed in our data. Under this account, the WM color information would have been carried along in the gradual decision formation throughout our task, without necessarily altering the sensory processing of the individual stream displays proper. In signal-detection terms^16^, the effect would be equivalent to a trial-by-trial shift in decision criterion to categorize a stream as blue (or red). In cognitive terms, the respective response category would be preactivated and chosen more liberally. From either perspective, the bias we observed would emerge at the level of decisional evaluation and -categorization.

An explanation in terms of category-level interference can also account for the observation that the bias in color reports was bidirectional. Memory reports were biased by the information in the intervening decision stream, including from late displays that arrived during presumably unprioritized WM storage. More generally, however, memory reports were attracted to the decisional evaluation result (i.e., “blue” or “red”), even regardless of whichever information was physically presented. Given the categorical nature of the (false) memory reports, we assume that they likewise reflected high-level conceptual interference (and/or -confusion), rather than the subtle alterations of a sensory memory trace^cf. 11,14^. Together, the bidirectional bias of WM- and decision reports may have arisen at a post-perceptual level of processing, independent from low-level sensory-mnemonic crosstalk.

Our findings integrate well with a view that WM contents can be flexibly maintained in different formats according to momentary task requirements^50–53^. Here, during a challenging visuospatial integration task, the concurrent color information may have been transferred to a more asensory (e.g., abstract-categorical, or verbal) format which avoids low-level perceptual interference^53^, but which may interact with task aspects that rely on similar levels of abstraction, such as decisional categorization along a shared feature dimension. While the present choice biases appear suboptimal in the context of our dual-task paradigm, the findings may connect to literatures on the role of trial history^17,54–57^, and of past experience more generally, in adaptive decision making. Prominent models, often in a Bayesian tradition, have characterized how prior experience shapes current percepts, and vice versa, how prior distributions are continuously updated through new experience^58–60^. Observing similar dynamics here in a dual task, we may speculate that the mechanisms behind concurrent WM storage may overlap with those that inject context information in adaptive decisions (for related discussion, see^61^), in exchange with longer-term memory and -knowledge^5,6,62^.

Neuroscientific work has offered a range of accounts how visual WM information might be stored throughout concurrent tasks. One set of recent neuroimaging studies has suggested an inverted coding scheme, where memories stored for current and prospective goals are encoded in opposite neural patterns^63–65^. While it yet remains to be shown if and how the neural representational quality of stored information impacts on concurrent perception, we found no evidence for inversion or suppression in spatial weighting patterns during prolonged storage for prospective use. We exclusively observed attractive biases. However, our psychometric method might not have been sensitive enough to detect more subtle, potentially inverse effects on later displays. Another recent functional imaging study suggested that unattended storage may rely on low-resolution representations in fronto-parietal areas, whereas attended storage additionally recruits high-precision representations in early visual cortex^52^. Our results may integrate with this view, although our design did not include a control condition to quantify loss of WM precision that is attributable to accessory storage in particular. How the interactions reported here relate to the neural representational nature of mnemonic- and perceptual information remains a question for future work using e.g., fMRI- and/or M/EEG recordings.

To conclude, our report indicates both sensory-perceptual and decisional processing stages as a potential locus of crosstalk between perception and concurrent WM. The findings complement and extend previous work on WM-perception interactions and support a view that concurrent WM storage may flexibly adapt to momentary task demands. We hope that the new approaches and methodologies introduced here will also prove instrumental in the continuing search for the neural substrates of WM.

## Methods

### Participants

In total n=80 healthy volunteers (43 female, 37 male; age 26.61 +/- 4.35) participated with written informed consent. Participants who failed to perform above chance level in WM color- (n=7) and/or location recall (n=2) were excluded from analysis. We further excluded n=3 participants whose choices in the decision task were not robustly driven by the blue-red dot count in the stream displays (p<0.001, logistic regression of choice). Results are reported for the remaining n=68 participants (36 female, 32 male).

### Stimuli and Task

Decision task. On each trial, participants viewed a stream of six circular displays (cf. Fig. 1, *middle;* outer circle diameter 10.6° visual angle). Each display contained 20 circular dots (diameter 0.3°), a varying number of which was colored blue, the others red. The difference in the number of blue – red dots in each display was randomly drawn from a normal distribution that was truncated to one standard deviation (SD=10). The mean of the distribution was randomly varied across trials to be either −4 or +4, however, due to an error in the presentation code it was consistently shifted by 1 dot towards red (in half of the subjects) or blue (in the remaining subjects). As this shift was small and constant in all trials and conditions of interest, we consider it inconsequential for the reported results. Control analyses confirmed that the overall color weighting functions in the two subgroups were virtually identical. The spatial positions of dots in each display were randomly assigned (independent of color) and uniformly distributed across the display area, with the restrictions that no dots overlapped and that the minimum distance to the outer border of the display area was 0.3°. Each display was presented for 0.2s, followed by a blank period (empty display area) of 0.1s. The outer circle and a central fixation cross remained on screen for the entire trial.

To foster participants’ motivation in the decision task, the red-blue comparison was instructed in an incentivized choice frame. One of the colors (red/blue, counterbalanced across participants) was designated as gain-, the other as loss color. Participants were instructed to accept the stream if it contained more gains, and to reject it otherwise, via left-hand button press (key “C” or “X”, respectively). After choice, full informative feedback was displayed, and accepted gains were credited to the participant’s running bonus balance. Feedback was based on the mean of the stimulus set (see above). In half of the participants, the task was framed as a gamble where the outcome (in terms of both feedback and bonus) was based on that of a randomly drawn display from the just-presented stream.

WM task. WM trials started with presentation of a single dot (diameter 0.3°; cf. Fig. 1 *left*) for 0.5 s, the color (red/blue, randomly varied across trials) and spatial location of which were to-be-remembered for later recall. The location of the WM sample varied randomly across trials but was restricted to a circular path of 3.8° radius around fixation. In WM interference blocks (Fig. 1a), the WM sample was followed after a 0.2s delay^46^ by the decision task. In Control 2 blocks (Fig 1b, lower), the WM sample was immediately followed by WM recall. On WM recall, a white dot appeared at the center of the display area and participants were asked to move it to the remembered location using arrow keys, and to toggle its color (red/blue) with key “0” on the numpad, all using the right hand. Participants were free to make adjustments and corrections (of both color and location) for as long as they wished before submitting their result by pressing the enter key. Thereupon, feedback of both color- and location accuracy (transformed into a percentage correct score) was displayed. Color- and location accuracy were combined into a bonus score that was surcharged on participants’ running bonus balance.

### Design and Procedure

Each participant performed 3 consecutive blocks of 80 trials in the critical WM interference condition, where the decision task was presented after WM encoding and before WM recall (Fig. 1a). Half of the participants additionally performed 3 blocks in control condition 1, in which the WM task elements were omitted (Fig. 1b, upper). The remaining participants performed 3 blocks in control condition 2, in which the WM- and decision task elements were reordered so that the two tasks were not concurrent (Fig. 1b, lower). The ordering of WM interference- and control blocks was counterbalanced across participants. All between-subject assignments (control 1/2, gain/loss color, choice framing, task order) were crossed to be orthogonal.

Participants where pre-experimentally instructed to give equal priority to the WM- and decision tasks, and that performance in both task components would be combined in the final bonus score. Participants were seated at approximately 57 cm viewing distance from a 24’’ TN display (BENQ XL2430, 531.36mm x 298.89mm viewing area, 144Hz refresh rate, 1920×1080 pixels resolution). A chinrest (SR Research) was used to minimize changes in viewing distance and head posture. Participants were instructed to fixate a centrally presented crosshair (10 x 10 pixels, 2 pixels linewidth) and to avoid eye movements during all task stages. After each block, summary performance feedback was provided, and participants were free to take a short break before continuing with the next block. Upon completion of the experiment, the bonus score balance was converted into a small monetary amount (2-5 Euro, depending on performance) and surcharged on the standard reimbursement for participation.

### Spatiotemporal weighting analyses

For spatial weighting analysis, the stream displays were reconstructed offline as 401×401 pixel circular pseudo-color maps (blue: 1, black: 0, red: −1) and smoothed with a 20×20 pixel Gaussian kernel. Spatial decision weight was estimated at each pixel (x,y) using a logistic regression of choice (Eq. 1):

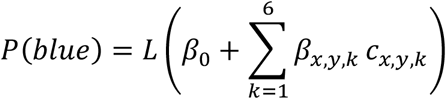

where *P*(*blue*) is the probability of choosing blue>red, *L* is the logistic function *y* = 1/(1 + *e^−x^*), and *c_x,y,k_* is the pseudo-color value at pixel coordinate x,y in stream display k. The coefficients *β_x,y,k_* form a spatiotemporal map (*M*) of decision weight in terms of psychometric slopes at each pixel x,y and each display position k (1-6), and *β*_0_ is the model’s constant term. We estimated *M* for each participant’s choice data and contrasted it with a spatially unbiased observer map *M\**. The unbiased map *M\** was estimated from the choice probabilities predicted by an individually fitted psychometric model (see *Psychometric model* below) which was uninformed by the displays’ spatial dot layout. Subtracting *M* − *M\** yields a map of regional over- (positive values) or underweighting (negative values). The maps were examined statistically using t-tests against zero with false-discovery-rate (FDR) correction^66^ for multiple comparisons in pixel space.

To test for concentration of decision weight at the concurrently maintained WM sample location, we recomputed *M* − *M\** from rotationally aligned stream displays. Specifically, we rotated the decision displays from each WM trial offline so that the WM sample location on any given trial was aligned at the same angular position (arbitrarily set to 45°). For instance, when the WM sample had been presented at 120°, the subsequent stream displays would be re-rotated by −75°. Trials with unusually inaccurate WM location recall (>40 pixels Euclidean displacement) were excluded from the main analysis. To statistically test for angular tuning, we partitioned the display area into 11 equal-sized pie segments and compared the average weight in the segment centered around the WM sample position to that in the remaining segments. The reported results pattern was robust to varying the number (width) of segments used in partitioning.

### Reverse correlation analysis of psychometric weight

Psychometric weighting functions (cf. Fig. 4) were derived by computing for each relative excess count of blue (vs red) dots, in each display position (1-6), the relative frequency with which the stream was subsequently judged blue (i.e., accepted when the gain color was blue or rejected when the gain color red). The slope of the resulting psychometric curve indicates the weight with which the blue-red dot count in a display impacted on final choice. In other words, the slope of the curve indicates the observer’s psychophysical *sensitivity* to this information. In contrast, an overall displacement of the function, e.g., in terms of a parallel shift away from an ideal observer function (which is point-symmetric at p=0.5) reflects an additive response bias towards one of the two choice categories, regardless of the stream’s physical appearance.

### Psychometric model

For quantitative analysis of choice bias and -sensitivity, we used a simple logistic choice model of the form (Eq. 2):

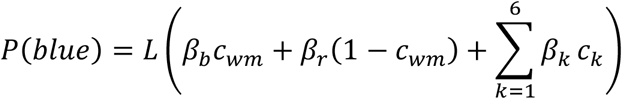

where *P*(*blue*) is the probability of choosing blue>red, *L* is the logistic function, *c_wm_* is a dummy variable coding the color of the WM sample (1: blue, 0: red), and *c_k_* is the color composition of stream display *k* in terms of the excess count of blue minus red dots (ranging from −10 to +10 relative to the mean of the stimulus set, see *Stimuli and Task*). The coefficients *β_k_* reflect the psychophysical sensitivity (slope) with which the color composition of a stream display is weighed in choice, whereas coefficients *β_b_* and *β_r_* reflect the strength of an additive choice bias associated with maintaining a red or blue WM sample, respectively. In control trials without a concurrent WM item, a single intercept (*β*_0_) was used instead.

### Statistical procedures

We used conventional parametric procedures (t-tests and ANOVAs) as detailed in *Results*. Greenhouse-Geisser corrected degrees of freedom were used where appropriate. Supplementary Bayesian statistics were computed using the Bayes factor toolbox (https://github.com/klabhub/bayesFactor)

### Ethics statement

The study was approved by the ethics commission of the Max Planck Institute for Human Development and was conducted in accordance with the Human Subjects Guidelines of the Declaration of Helsinki.

## Data availability

The research data supporting the findings will be made available upon publication through the institutional git repository:

## Code availability

The experimental and analysis code will be made available upon publication through the institutional git repository:

## Acknowledgement

We thank Thomas Christophel, Martin Rolfs, Martin Spitzer, and Juan Linde-Domingo for helpful discussions, Jann Wäscher for help with data acquisition, and Agnessa Karapetian for assistance. Conception of the work was supported by a grant from the German Research Foundation (DFG) to BS (SP 1510/1-2); ZK was supported by the Berlin School of Mind and Brain.

